# Compounds without borders: a novel paradigm for quantifying complex odors and responses to scent-pollution in bumblebees

**DOI:** 10.1101/684134

**Authors:** Jordanna D. H. Sprayberry

## Abstract

Bumblebees are critical pollinators whose populations have been experiencing troubling declines over the past several decades. Successful foraging improves colony fitness, thus understanding how anthropogenic influences modulate foraging behavior may aid conservation efforts. Odor pollution can have negative impacts on bumble- and honey-bees foraging behavior. However, given the vast array of potential scent contaminants, individually testing pollutants is an ineffective approach. The ability to quantitatively measure how much scent-pollution of a floral-odor bumblebees can tolerate would represent a paradigm shift in odor-pollution studies. Current statistical methods for analyzing complex odors have poor predictive power because statistically-derived odor-spaces are rewritten when new odors are added. This study presents an alternative method of analyzing complex odor blends based on the encoding properties of insect olfactory systems. This “Compounds Without Borders” (CWB) method vectorizes odors in a multidimensional space representing relevant functional group and carbon characteristics of their component odorants. A single vector can be built for any scent, which allows the angular distance between any two odors to be calculated – including a learned odor and its polluted counterpart. Data presented here indicate that CWB-angles are capable of both describing and predicting bumblebee odor-discrimination behavior: odor pairs with angular distances in the 20-29° range are generalized, while odor pairs over 30 degrees are differentiated. The neurophysiological properties underlying CWB-vectorization of odors are not unique to bumblebees; CWB-angle analysis of published data on a hawkmoth supports the idea that this method may have broader applications.

## INTRODUCTION

Bumblebees are prolific pollinators in both natural and agricultural ecosystems [1,2]; which makes the decades long decline in bumblebee populations [3] particularly alarming. Given that the foraging success of workers can be directly linked to reproductive output of a colony[4], understanding both general mechanisms of foraging and how anthropogenic environments impact foraging is directly relevant to conservation efforts. Flowers provide multiple sensory advertisements to pollinators; such as shape, color, and scent[5–7]. Recent computational work indicates that odor is consistently available to searching bumblebees[8] and lab-based experiments indicate that bumblebees are capable of using odor information alone to locate floral resources [9,10]. Floral-scent is likely an important sensory cue for bumblebee foragers; unfortunately anthropogenic activity has modified their olfactory landscape in urban, suburban and agricultural ecosystems [11–15]. Air pollution can react with floral odorants resulting in modified floral-odor blend structure and reduced ability of honeybees to recognize a learned odor [12,13]. Agrochemicals can have strong odor signatures which have been shown to modulate bumblebee foraging behavior [10]. Given the vast quantity of potential odor pollutants in human habitats, it is unrealistic to approach this problem through testing them one by one. However, the ability of foraging bumblebees to successfully recognize and locate scattered resources may be crucial to species’survival. Therefore, an effective method to predict the likelihood that a given odor pollutant will disrupt foraging behavior is needed. Current methods of describing the relative similarity (or dissimilarity) of complex odor blends rely on statistical analysis, typically principal components analysis (PCA) [16–19]. This has little predictive power as the statistically-based odor space is recreated whenever a new odor is introduced. This study proposes a multidimensional odor-space based on the current understanding of how odorants are encoded by insect-olfactory systems; tracking their functional group and carbon characteristics [16,18,20-25]. While there is little experimental evidence on olfactory processing from bumblebees [9,26], given the homology across insect olfactory systems it is reasonable to hypothesize that bumblebees have commonalities with other species [21,27]. Studies on olfactory receptor neurons (ORNs) indicate that many ORNs respond to multiple molecules sharing functional group characteristics [20,24]. Seminal work on olfactory processing in the honeybee antennal lobe demonstrated that carbon chain length and functional group are reliably encoded [18]. Their findings were supported by later work on responses to complex floral odors in hawkmoths showing that scents whose components have the same functional group elicited similar responses from the antennal lobe [16]. Moreover, recent work on the antennal lobe tracts (ALT) leaving the antennal lobe in honeybees show that the mALT carries information about functional group, while the lALT appears to encode carbon chain length [25]. The odor space presented here builds multidimensional vectors based on the distribution of functional group and carbon characteristics within a complex odor blend because they should have a meaningful relationship to odor-driven behaviors. The angular distance between vectors of two complex odors can serve as a measurement of their similarity (or lack there of). Unlike PCA these dimensions exist independently of the odors they characterize, thus the odor space is constant regardless of how many odors are being analyzed. We tested the efficacy of this “Compounds Without Borders” (CWB) odor-characterization method with an associative odor learning and discrimination paradigm (free-moving proboscis extension reflex, FMPER[28]). Once the viability of the CWB-method was established, we applied it predictively – successfully hypothesizing bumblebee response to a novel set of odor-discrimination tasks. This represents both a new mechanism of computationally characterizing complex, ecologically-relevant odor blends and provides a new tool to predict which types of odor pollution are more likely to disrupt odor recognition.

## RESULTS

### Using ‘Compounds Without Borders’ to calculate angular distances between odor-pairs provides the ability to measure similarity with a single quantitative variable

Characterization of odor-blends entailed identifying component odorants, calculating their normalized peak areas, and determining their dimensional characteristics based upon their respective carbon chain length (CCL), cyclic carbon count (CCC), and functional groups (FG) (Fig. 1, Table S1). The vector for the odor-blend was calculated by summing the area for all component odorants within each dimension. This ‘Compounds Without Borders’ (CWB) method of vector construction allows the calculation of angular distances between any two odors regardless of their underlying complexity, quantifying their relative similarity (or lack thereof) with a single variable. The CWB-angles between the three primary odors used to explore the viability of this method are shown in Fig. 1C: Lily of the valley (LoV) is closer in structure to honeysuckle (HS) than juniper berry (JB), but JB is closer to LoV than HS (Table 1).

**Table 1.** Stimulus characteristics and response details for all discrimination tasks. The ‘random’ distribution assumes an equal probability of correct, incorrect, and no-choice responses, while ‘expected’ represents the theoretical response. C = correct, I = incorrect, and NC = no choice.

| Associative Odor | Comparison Odor | Angle | p (vs random) | p (vs expected) | % C/I/NC | n |
| --- | --- | --- | --- | --- | --- | --- |
| LoV | MO | n/a | 0.0006 | 0.08 | 66.7/15.2/18.2 | 33 |
| LoV | LoV | 0 | n/a | 0.003 | 27.8/22.2/50 | 15 |
| LoV | 3 LoV: 1 JB | 14.4 | n/a | 0.05 | 47.6/0/52.4 | 21 |
| LoV | 1 LoV: 1 HS | 26.8 | n/a | 0.05 | 33.3/6.7/60 | 15 |
| LoV | 1 LoV: 1 JB | 30.5 | n/a | 0.18 | 53.3/20/26.7 | 15 |
| LoV | HS | 56.5 | n/a | 0.14 | 60/20/20 | 15 |
| LoV | JB | 58.9 | n/a | 0.32 | 61.9/0/38.1 | 21 |
| JB | MO | n/a | 0.002 | 0.13 | 71.4/14.3/14.3 | 21 |
| JB | JB | 0 | n/a | 0.0009 | 22.2/22.2/55.6 | 18 |
| JB | 3 JB: 1 LoV | 13.0 | n/a | 0.09 | 56.7/0/43.3 | 30 |
| JB | JB: 1 LoV | 28.5 | n/a | 0.03 | 33.3/20/46.7 | 15 |
| JB | 1 JB: 1 HS | 31.7 | n/a | 0.68 | 53.3/13.3/33.3 | 15 |
| JB | LoV | 58.9 | n/a | 0.43 | 70.6/0/29.4 | 17 |
| JB | HS | 69.8 | n/a | 0.50 | 64.7/0/35.3 | 17 |
| Crd | MO | n/a | 0.03 | 0.30 | 50/13.3/36.7 | 30 |
| Crd | 1 Crd: 1 Brg | 24.8 | n/a | 0.02 | 38.9/0/61.1 | 18 |
| Crd | Brg | 56.7 | n/a | 0.23 | 48.6/8.1/43.2 | 37 |

**Fig. 1.**
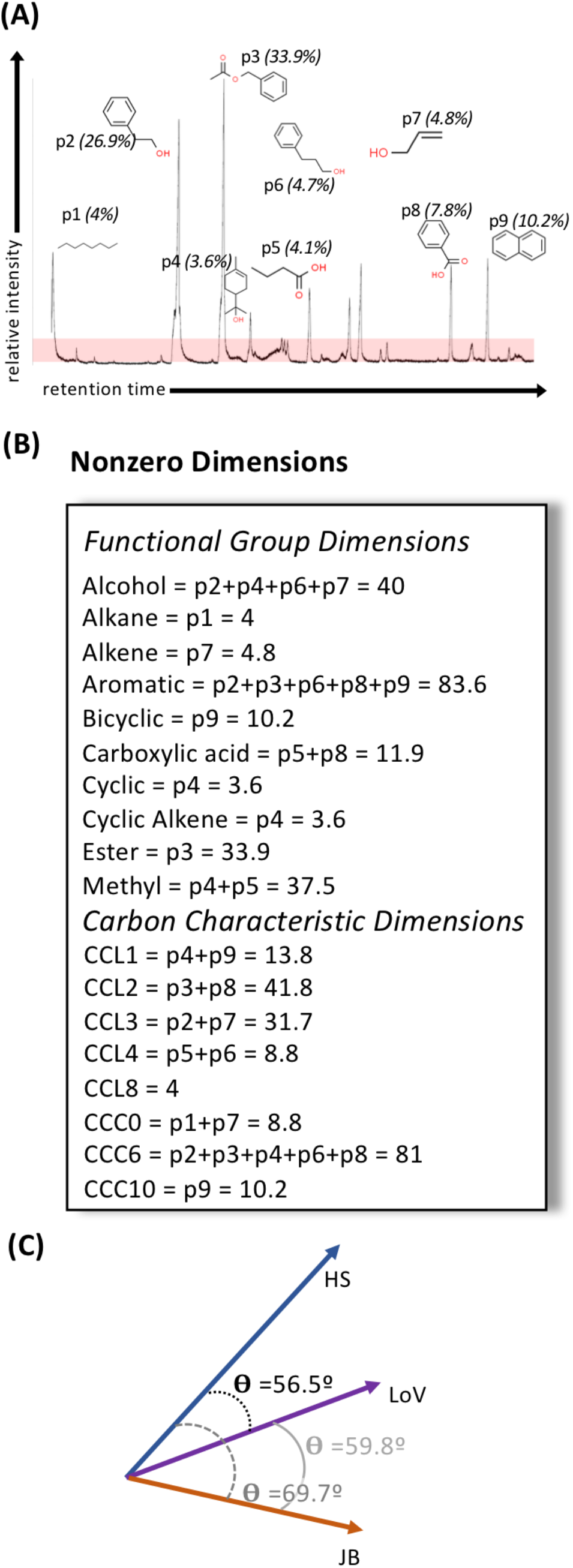
CWB-vectorization of honeysuckle odor.(A) The GC trace for honeysuckle essential oil, with peaks above 10x noise identified. The nine peaks were: p1= octane, p2=phenythyl alcohol, p3=acetic acid phenylmethyl ester, p4= α-terpineol, p5=butanoic acid, p6= benzenepropanol, p7=2-propen-1-ol, p8=benzoic acid, p9=napthalene. The structure of and relative area under the curve for all identified peaks is noted. (B) Each peak was characterized based on its functional groups, carbon chain length (CCL) and number of carbons in cyclic structures (CCC). A dimension receives a non-zero value if one or more odorants within an odor blend possess that characteristic. The relative peak area for all peaks with a particular characteristic are summed to determine the total power for that dimension. This method does not treat individual odorant compounds as discrete entities; rather it acknowledges that multiple different compounds may have the ability to bind to a single species of odorant receptor and that a single compound may bind multiple species of odorant receptor [20,24]. Thus by removing the boundary around individual molecules, this “Compounds Without Borders” method may more accurately model stimulation of the olfactory system. (C) The angular distance between any two CWB-vectors can be calculated, allowing the similarity or dissimilarity of two odors to be represented with a single quantitative variable.

### FMPER is an effective method for measuring odor learning

We modified the free-moving PER method presented by Muth et al. [28] to allow for odor stimulation (Fig. 2A). Bumblebees participating in FMPER had four potential outcomes: 1. they could fail to complete four conditioning trials and be dropped from analysis; 2. they could choose correctly, extending their proboscis in response to the AO-strip; 3. they could choose incorrectly, extending their proboscis in response to the CO-strip; or 4. they could not choose.

**Fig. 2.**
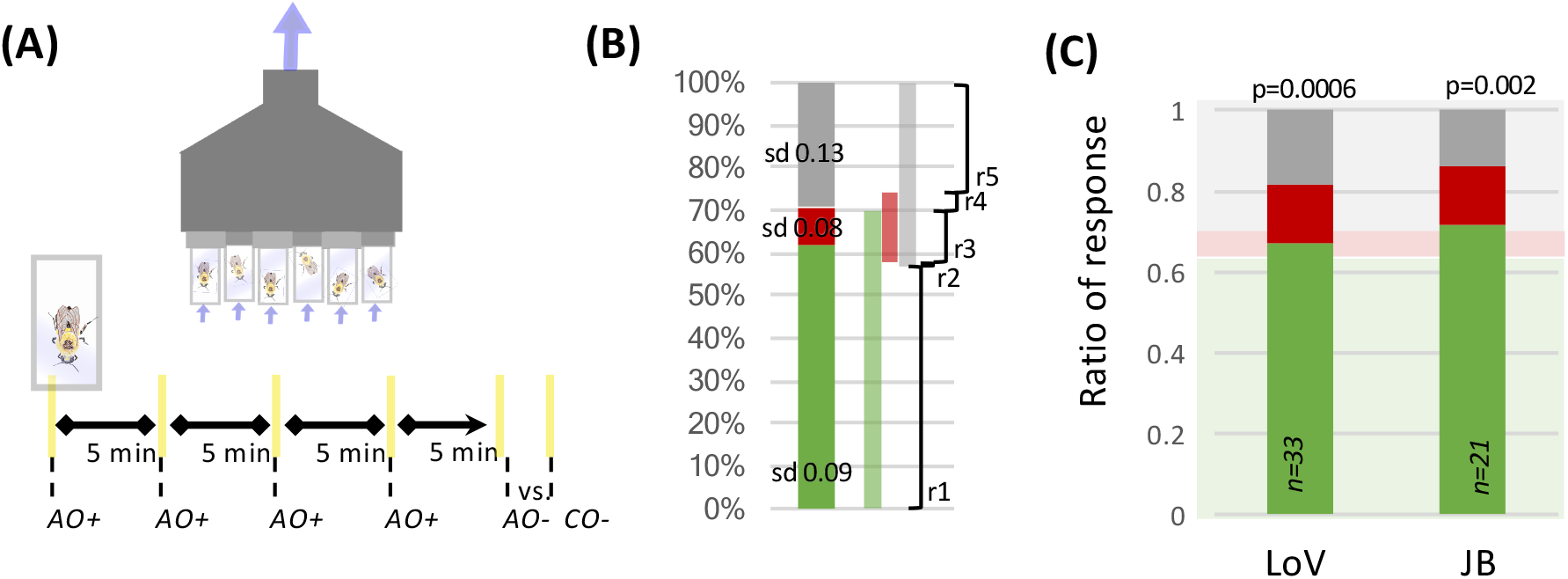
(A) The ventilating array held 6 bees in screen-backed vials, bringing air in through two lid-holes and out the back. A sucrose reward was delivered on yellow AO-scented plastic strips during the four conditioning trials. The testing trial presented bees with two unrewarding strips (AO vs CO) and recorded bumblebee responses as correct, incorrect, or no-choice. (B) The mean and standard deviation for correct (green), incorrect (red), and no-choice (grey) response distributions for 7 different associative odor (AO) versus unscented mineral oil tests, utilizing a total of 177 bees, were used to statistically model the theoretical response distribution for an easy odor choice in this FMPER paradigm. A random number generator determined what range of response probabilities a silica-bee fell into: range 1 (r1) had 100% probability of being assigned “correct”, range 2 had an equal probability of being correct or no-choice, range 3 had an equal probability of being correct, incorrect, or no-choice; range 4 had an equal probability of being incorrect or no-choice, and range 5 had a 100% probability of being no-choice. This model returned a response probability-distribution of 63% correct, 6% incorrect, and 31% no-choice. (C) The results of AO versus mineral oil tests for lily of the valley (LoV) and juniper berry (JB). The background green, red and grey boxes represent the expected response distribution. The p-values and sample sizes are noted for each test.

Bumblebees were classified as “no-choice” if they successfully completed four conditioning trials but did not choose after interacting with the test strips three times. 446 individual bees from 10 *Bombus impatiens* colonies that were tested in these FMPER experiments. Of these 446, a total of 89 individuals (20%) were excluded from analysis, leaving 357 for analysis: 27 (6%) did not complete all four training trails; 17 (4%) were removed because of experimenter or equipment error; and 45 (10%) were from individual experiments resulting entirely in ‘no-choice’ (unfiltered data available for download).

Bumblebees demonstrated an ability to discriminate between an associative odor (AO) and unscented mineral oil (Fig. 2C): those associated to LoV yielded 66.7% correct (p=0.0006; exact test against random distribution) and association to juniper berry (JB) showed 71.4% (p=0.002). These data were combined with results from tests of 5 additional types of associative odors (see Table S2) to statistically-model an expected response distribution for the FMPER assay when bees were given an easy odor discrimination task (a perceivable AO versus unscented-mineral oil). The model returned a distribution of 63% correct, 6% incorrect, and 31% no choice (Fig. 2B). Responses to more challenging odor discrimination tasks are tested against this distribution with the interpretation that a statistical comparison yielding a small p-value indicates that the two tested odors are not easily discriminated, and likely being treated similarly – a phenomenon referred to as generalization [29].

### FMPER responses indicate that CWB-angles can identify a threshold for discrimination

Bumblebees associated to LoV were tested against: LoV, HS, JB and a range of corresponding blends (creating odor pollution of the AO) that resulted in an angular range of 0-58.9º. Bumblebees associated to JB were likewise tested against JB, HS, LoV and a range of corresponding blends that resulted in an angular range of 0-69.8º. In both AO conditions bumblebee response distributions to the 0º task returned a very low p-value when compared to the expected distribution (LoV vs LoV, p=0.003, n=15; JB vs JB, p=0.0009, n=18). Interestingly, both these tests showed a larger than expected percent no-choice: 50% for LoV-associated bumblebees and 55.6% for JB-associated (Fig. 4). In both AO conditions, responses to discrimination tasks with an angular distance larger than 30º had distributions that were similar to the expected response, with p values ranging from 0.14-0.68 (Fig. 3, Table 1), with high percent correct (>50%); indicating that the AO and CO were easily discriminated. Reponses to angles below the 30º threshold were more complicated. Bumblebees given AO vs CO tests with distances in the 20-29º range showed response distributions that were similar to the 0º tests: low percentages correct (< 40%), higher percentages of no-response (46.7-61.1%) and small p-values (Fig 3, Table 1); indicating that the CO was likely generalized to the AO. However, they did have higher percent correct than incorrect. Responses to angular separations of 10-15º show an increase in percent correct and a complete absence of incorrect choices. For the JB vs 3 JB: 1 LoV test (θ=13º), with 56.7% correct, the response distribution is not clearly different than expected (p=0.09). The LoV vs 3 LoV: 1 JB test (θ=14.4º) had 47.6% correct and did have a distinctly different response distribution than expected (p=0.05). Despite the statistical differences in how these two tests relate to the expected (easy odor choice) distribution, both demonstrated a high number of no-choice responses (43.3% for AO=JB and 52.4% for AO=LoV).

**Fig. 3.**
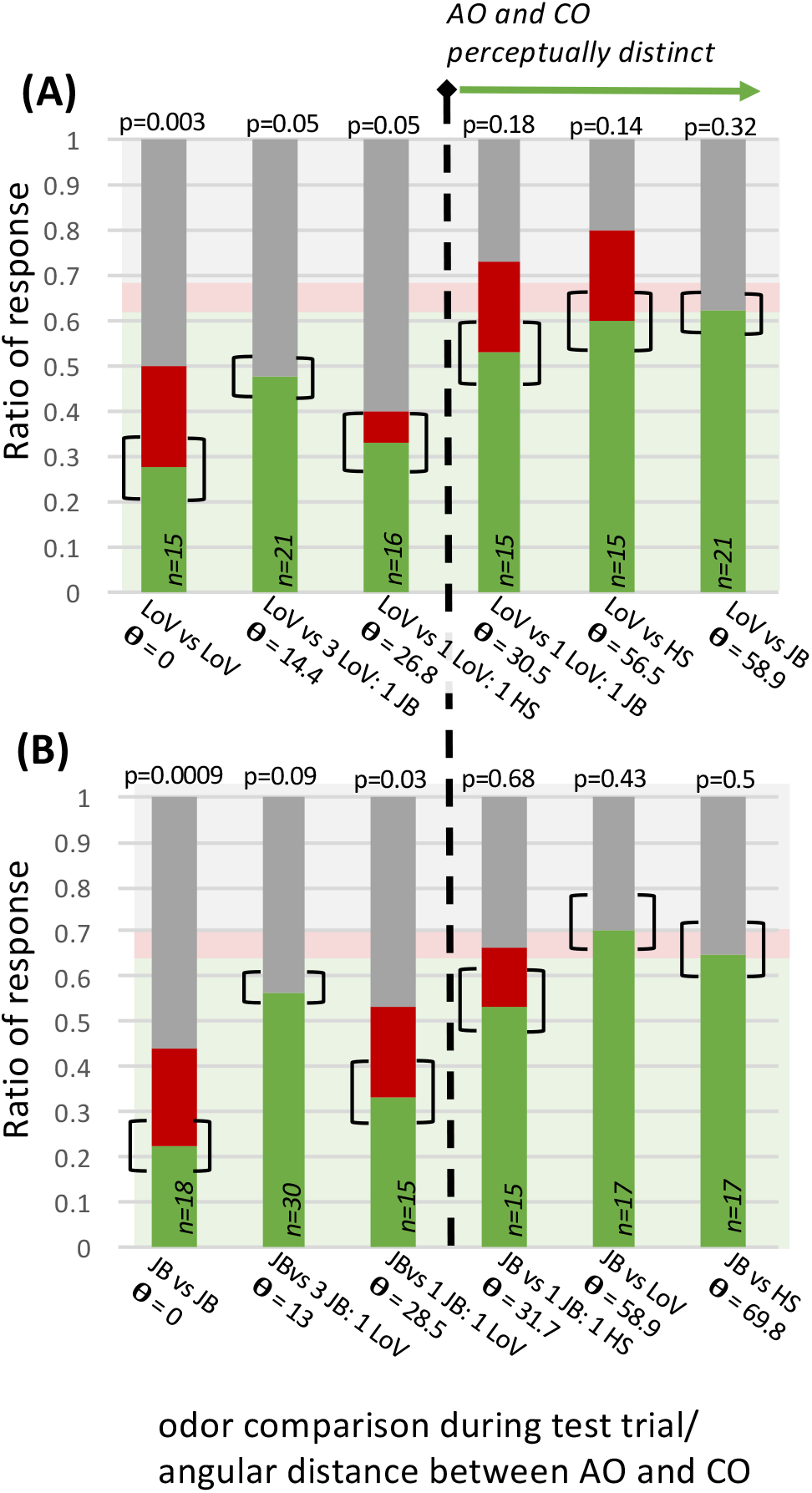
The response distributions for bumblebees associated to lily of the valley (A) and juniper berry (B) to discrimination tasks against contrasting odor (CO) stimuli with increasing angular distances to the associative odor (AO). Green portions of bars represent correct responses, red represent incorrect, and grey represent no choice. The transparent green, red and grey boxes in the background represent the expected response distribution. The p-values for the statistical comparisons with this distribution and sample sizes are noted for each task. All odors above 30º appear to be easily discriminated, as indicated with the vertical dashed line. The brackets around the correct response ratios represent the difference one bee makes (i.e. if one more or one less bee chose correctly).

**Fig. 4.**
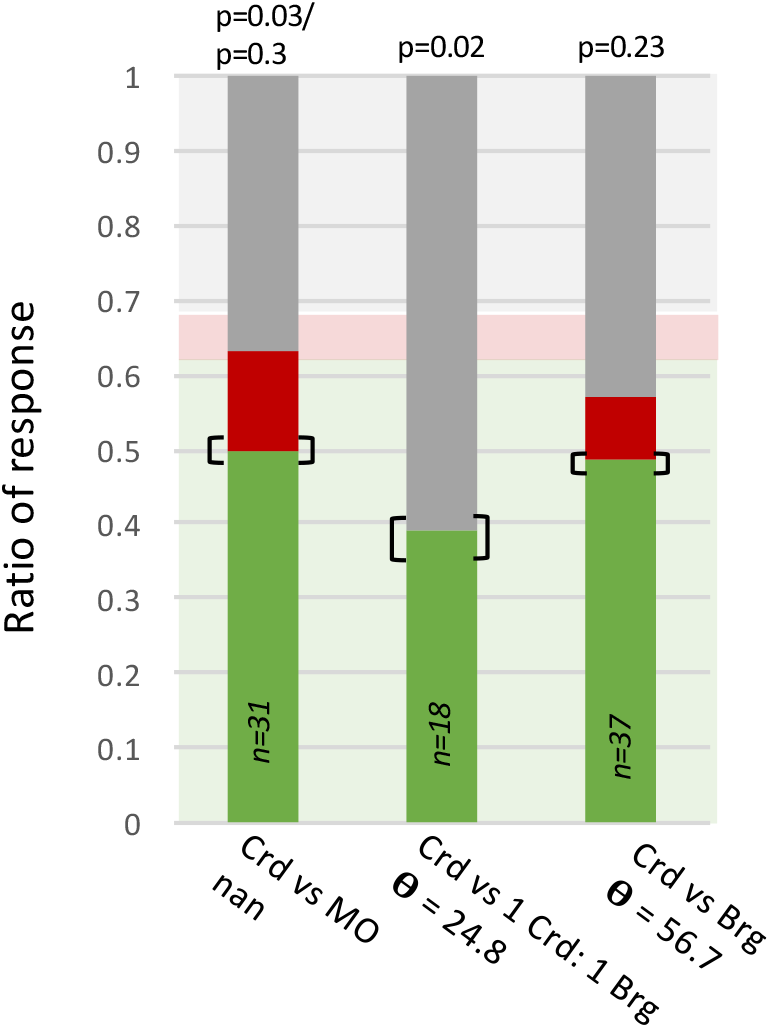
The response distributions for odor discrimination tasks testing the putative 30º discrimination threshold. Bumblebees were associated to cardamom (Crd) essential oil and tested against: mineral oil (confirming bumblebees are capable of learning Crd), a subthreshold CO (θ= 25.1º, Crd vs 1 Crd: 1 bergamot (Brg)), and a suprathreshold CO ((θ= 57º, Crd vs Brg). Green portions of bars represent correct responses, red represent incorrect, and grey represent no choice. The transparent green, red and grey boxes in the background represent the expected response distribution. The brackets around the correct response ratios represent the difference a change in one bee’s response would make.

### The CWB-method is capable of predicting bumblebee-responses in a FMPER task

The principle goal in developing the CWB-method was to facilitate characterization of odors in a predictive manner. Thus when preliminary data analysis indicated a putative threshold of discrimination, CWB-vectors were used to select two additional AO vs CO tasks: one with a subthreshold angular distance between the AO and CO, and one over threshold. The subthreshold task should yield a response distribution that differs from expected, while the suprathreshold task should be similar to the expected distribution. Using cardamom essential oil as the AO, the sub-threshold CO was 1 cardmamom: 1 bergamot (θ= 24.8º), and the suprathreshold CO was bergamot essential oil (θ= 56.7º). Indeed, the distribution for the 56.7º task appears to match the expected easy odor choice distribution (p=0.23), while the distribution for the 24.8º task differs (p=0.02) (Fig. 4)– thus the tested responses matched the apriori predictions.

### Scent contamination can render a learned odor unrecognizable

The threshold for discrimination cuts right through the cluster of COs that were constructed by blending a contaminating scent with the AO (Fig. 5). In these cases, bumblebees were clearly able to discriminate between the uncontaminated AO and its polluted CO, indicating that the polluted version of the learned odor was no longer being treated as the AO itself. However, bumblebees appear to have tolerance for scent contamination in the 20-29º range. Bumblebees tested with COs from this range showed a decreased percent correct and a departure from the expected response distribution, indicating the contaminated scent was difficult to discriminate from the original and likely generalized.

**Fig. 5.**
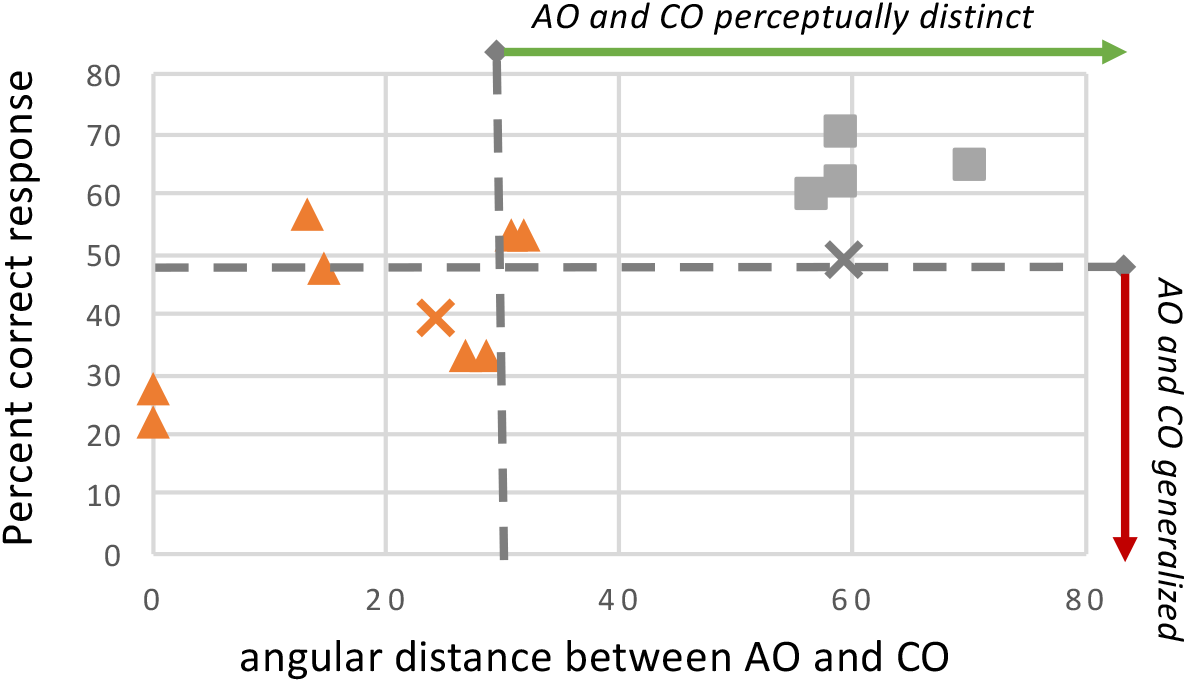
The percent correct data from all discrimination tasks replotted against the angular distance between the AO and CO. Data are differentiated by whether the CO was a blend of the AO with a polluting odor (▲) or if the CO was a distinctly different scent (▪). The predictive data (AO=cardamom) are represented with x’s: **X** for AO-blended CO and **X** for the distinctly different CO. Tests with % correct values below the horizontal dashed line consistently had small p-values when compared to the expected response distribution. The vertical dashed line at 30º represents the threshold above which bumblebees can easily discriminate between the AO and the CO.

## DISCUSSION

### Compounds without borders effectively describes olfactory discrimination behavior in an associative learning context

While the role of odor signals in plant-pollinator relationships is not completely understood, numerous studies indicate that odor is an important sensory modality in bumblebee foraging [9,30-32]. Given the complex olfactory landscapes that pollinators operate within [33], discrimination thresholds could have consequences for foraging efficiency. In an environment where target flowers (i.e. those a pollinator has coevolved with) have similar scents [16], foragers with a larger discrimination threshold angle and a generous generalization range could maximize feeding opportunities. Under the baseline conditions in this study (absolute conditioning and discrimination tests without an odor background) bumblebees showed a generalization range from 20-29°with a discrimination threshold of 30°. These angular ranges were effective descriptors of bumblebee behavior when associated to two different complex odor blends, lily of the valley and juniper berry, *and* served as an effective predictor of behavior for bumblebees associated to cardamom. However, the discrimination threshold described here does not likely represent a perceptual limitation. Work on ants has shown that previously generalized odor pairs can be discriminated after training with differential conditioning, implying that the behavioral-discrimination measured was contextual [29]. Indeed, the role of experience in generalization behavior has been demonstrated in multiple studies [34,35]. This behaviorally-demonstrated ability to dynamically shift discrimination thresholds in insects has been supported by work showing that learning can modulate neural activity within the antennal lobe [16,17]. Within the dataset presented here, responses to AO vs CO tasks with a 10-15° separation also support the idea that the 20-29°generalization range does not represent a perceptual limitation. Angles within this close range do not always appear to be generalized: bumblebee ability to correctly locate unpolluted JB when contrasted against a 3JB: 1 LoV blend (56.7%, 13°) is much higher than when tested against a 1JB: 1LoV blend (33.3%, 28.5°). Statistically, bumblebees associated to LoV and tested against 3LoV: 1JB (14.4°) did appear to generalize these two odors; yet their response distribution had 47.6% correct compared to the 31.3% of 1LoV: 1JB (14.4°), and showed no incorrect responses. This could imply that the lower threshold for generalization lies between 13 and 14.5°, or that there is a perceptually-distinct but behaviorally-ambiguous range where responses are less predictable. There are several potential ecological and ethological reasons why a range of ambiguity might exist. For example, recruitment pheromone activates foraging in bumblebee workers [36,37], and newly activated foragers are more likely to seek out floral odors brought into the hive by a returning worker [30]. Foraging pheromone would increase the power of eleven different CWB dimensions (Table S3), three of them with contributions from multiple components [30]. Depending on the volume released, the angular shift induced by pheromonal-‘pollution’ of floral odors brought in by returning foragers is likely to be small. Newly recruited foragers would want to generalize that small angular shift to the original floral blend, otherwise they risk not recognizing that floral resource. However, some small angular shifts might be disadvantageous to generalize. Recent work on volatile emission from microbial nectar communities has shown differences in their odors. Interestingly honeybees show differential attraction to microbial scents [19]. If the angular shift in floral odor induced by a microbial community is small, *and* if the added odor is indicating the presence of an undesirable microbe, than generalizing the microbe-polluted floral odor to the original would be disadvantageous. The ecological, ethological, neurophysiological, and perceptual drivers of behaviors to odor shifts in this angular range are fascinating topics for future study.

### Understanding generalization and discrimination behavior of bumblebees could help conservation efforts to reduce the impact of odor pollution

As of June 2019 Home Depot’s website listed 333 products under plant care geared for disease control and fertilization. At least one prior study has shown that the scent of a consumer lawn product is capable of modifying bumblebee foraging behavior in the lab [10] - but piecewise testing of 333 products is an intractable amount of work for the academic community and agrochemical producers are unlikely to take up the cause. Agrochemical odor pollution is not confined to consumer products; commercial agrochemicals may also be problematic [10]. Using the CWB-method to calculate angular shifts due to agrochemical scent-pollution could identify products that shift odors into a ‘zone of concern’, namely outside the identified generalization zone. This could be a remarkably useful tool for predicting which products are likely to disrupt bumblebee recognition of a learned floral resource. Likewise, this may provide a strategy for encouraging bumblebees to avoid resources recently treated with insecticides. However, all real-world ecological applications will require field testing.

Unfortunately, agrochemicals are not the sole source of odor pollutants that pollinators contend with – previous work has shown that air pollutants, such as diesel exhaust and ozone, react with floral odorants[11–15]. This reduces the distance that floral odors travel, which could have impacts on signal encounter by searching foragers [8,12]. In addition, the reaction of floral odorants with air pollutants changes the blend structure – in some cases pushing a learned odor far enough that honeybees no longer respond normally [13–15]. CWB-angles may provide a useful tool for investigating what levels of air pollution are likely to disrupt learned odor responses, and which still allow generalization to known scents.

### Broader impacts and limitations of the CWB-method

Due to the limited neurophysiological data from bumblebee olfactory-systems [9,26] the architecture of the CWB-dimensions was based on the encoding properties of other insect species [38], [18,39], [24,40]. Therefore, the CWB-method may be applicable to insects more broadly. Riffell et al. assembled a comprehensive data set in their seminal study on the coevolutionary and neurophysiological relationships between hawkmoths and hawkmoth-pollinated flowers, where they used PCA to show that evolved pollinator relationships have a stronger effect on floral odor structure than phylogeny, that innate neural responses to floral odors clustered based on the odors’ functional group attributes, and that a subset of floral odors were more innately attractive to hawkmoths[16]. To determine if CWB-angles are capable of recapitulating traditional PCA analysis, the CWB-angles between the first (randomly selected) hawkmoth-pollinated flower from their figure 1a, *Nicotiana suaveolens*, and all other tested odors were calculated from the odor structure data provided in their supplemental materials. Indeed, CWB-angles perfectly mirrored Riffell et al.’s analysis of scent structure (Fig. 6). Every hawkmoth-pollinated floral odor that clustered together in Riffell et al.’s PCA of odor structures has a CWB-angle of 40° or less, while every odor outside the PCA-cluster has an angle of 44° or higher. Moreover, with few exceptions, CWB-angles also correlate with their PCA analyses of neurophysiological and behavioral data (Fig. 6). This implies that analyzing odors with the CWB-method may be effective in some contexts where PCA is currently applied. However, this method has only been tested with complex odor blends. Odor-driven behaviors, such as pheromone tracking, based on smaller numbers of odorants may be less effectively characterized depending on the inter-dimensional overlap of the component compounds. While the work presented here provides proof of concept, its inter-species and ecological applications will require additional experimental validation.

**Fig. 6.**
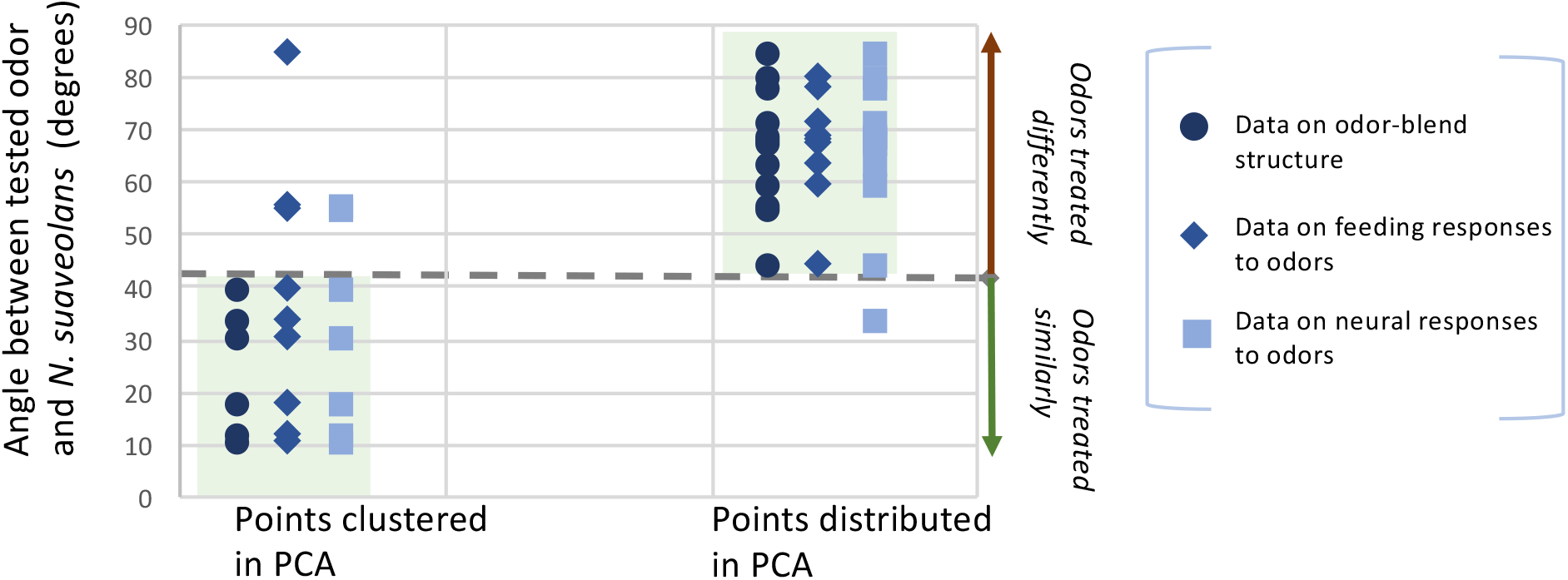
Comparison of CWB-angle analysis to PCA of data from Riffell et al. indicates CWB-angles are capable of recreating traditional approaches to complex odor-blend analysis. Riffell et al. used PCA determine relationships between odor-blend structures (●), as well as the relationship between odor and hawkmoths’ innate behavioral (♦) and neural (▪) responses. A hawkmoth-pollinated flower, *Nicotiana suaveolans*, from their data was randomly selected as the anchoring odor, and the CWB-angles to all other floral-odors they presented were calculated; these values are the y-axis. The left hand side shows odors/ responses that were clustered in their PCA, while the right hand side shows odors/ reponses that were outside their identified clusters. The transparent green boxes indicate areas where CWB-angles overlap with the PCA.

## METHODS

### Odor stimuli

We selected three essential oils (*New Directions Aromatics)* to serve as odor-blends for this study: lily of the valley, honeysuckle and juniper berry. Lily of the valley (LoV) has been successfully used in odor learning experiments utilizing the proboscis extension reflex (PER) [41], and the two additional odors were selected to provide a range of structural overlap with LoV. For further discussion of odor stimuli selection, see supplemental materials. Blending these three in varying ratios allowed construction of odor stimuli with varying ranges of odorant composition and distance (Table 1). These selected essential oils, as well as those from the predictive FMPER experiments, were sampled using Solid-Phase Microextraction (SPME) fibers and their composition analyzed with GCMS. For full details please see supplementary methods.

### CWB-vectorization of odor-blends and calculation of angular distances

Characterization of odor blends identified component odorants and their normalized peak areas. The dimensional signature of each odor-blend was then determined by the molecular structure of its component odorants based upon their respective carbon chain length (CCL), cyclic carbon count (CCC), and functional group (FG) characteristics (Fig. 1). From a dimensional perspective, the power for each dimension was calculated as the summed area of all peaks from molecules with that attribute (CCL, CCC, or FG). If no molecules within a given odor blend have that attribute, that dimension has a power of zero. From a peak/ molecule perspective, each odorant’s normalized area will be assigned to multiple dimensions; with a minimum of two (CCL and at least one FG) and no maximum (Fig. 1).

The CWB method represents odors as vectors in a 66-dimensional space. This vector representation allows us to calculate angular differences between odor blends, where given two odor vectors a and b the angle between them can be calculated as:

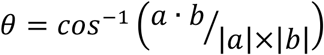

The calculated vectors and the odor blend classifications are available by request and the R code for calculating vectors and angles are available in Appendix S1.

### Associative testing of odor discrimination with FMPER

Testing the efficacy of the CWB-method of odor representation required a reliable method of assessing odor learning and discrimination. We modified the free-moving PER method presented by Muth et al. [28]to allow for odor stimulation (Fig. 2A). Healthy-, active-individual *Bombus impatiens* (from Kopert Biological) were selected from lab colonies, placed in screen backed vials, acclimated for two hours, and placed into the odor stimulation apparatus. The ventilating testing array drew air in through two small holes in the lid and out the back, with flow rates ranging from 0.1-0.3 m/s (VWR-21800-024 hot wire anemometer). During conditioning bees were offered a single drop of sucrose on a yellow strips cut from plastic folders, which had absorbent adhesive bandage tape (Cover Roll) placed on the back to hold associative odor stimuli (1 µL of essential oil). The plastic prevented the odor-solution from diffusing into the sugar solution on the tip of the strip, therefore the primary sensory encounter with odorants was through the olfactory rather than the gustatory system. Four conditioning trials were followed by an unrewarded test trial with five minute inter-trial intervals (Fig. 2A). Conditioning trials were not started until a bee successfully consumed sucrose. Test trials presented conditioned bees with two unrewarding strips-one scented with AO and the other with the CO. The complete protocol for these experiments can be found in Appendix S2. The contrasting odors are detailed in Table 1. Bumblebees participating in FMPER had four potential outcomes: 1. They could fail to complete four conditioning trials and be dropped from analysis; 2. They could choose correctly, extending their proboscis on the AO strip or while antennating the AO strip; 3. They could choose incorrectly, extending their proboscis on the CO strip or while antennating the CO strip; or 4. They could not choose. Bumblebees were classified as “no-choice” if they successfully completed four conditioning trials but when presented with two unrewarded strips during a test trial approached and investigated strips three times without choosing. On the rare occasion that all bees tested on a given day returned ‘no-choice, those data were excluded from the overall data set (45/446 bees). All FMPER data are available by request.

### Statistical analysis of FMPER results

Statistical analyses of FMPER results asked two questions: 1) does FMPER provide a reasonable measure of associative olfactory learning; and 2) are two odors difficult to discriminate from each other?

To determine if FMPER tests associative olfactory learning, bumblebees responses to a discrimination task of the AO from unscented mineral oil (MO) were tested against random chance with an exact test of goodness of fit [42].

Given that FMPER did indeed provide a reasonable measure of odor-learning (Fig. 3), the response-distribution of bumblebees tested with an AO vs MO should represent a simple odor discrimination task. We used data from the three AO vs MO experiments in this study, as well as four aditional AO vs MO experiments from a separate methods study (Edwards et al. *in prep*) to establish the mean correct, incorrect and no-choice responses made by bumblebees in this simple task (Table S2). These data were used to statistically model the a theoretical response distribution of how bumblebees respond to an easy odor discrimination task in the FMPER assay (Fig. 2B). Full details on this model are available in the supplemental materials (code available in Appendix S3). In order to answer the question, ‘were the two tested odors (the AO and the CO) difficult to discriminate?’ we tested response distributions for each test against the theoretical distribution (63% correct, 6% incorrect, and 31% no choice) with an exact test of goodness-of-fit using a log liklihood ratio method for calculating p values. Therefore the null hypothesis (accepted if p values are large) would be that the two odors are in fact easily discriminated. Rejection of the null hypothesis would indicate that the odors are not easily discriminated, or are generalized. For readers wanting to assess traditional ‘significance’ of p-values of these analyses may consider using an alpha level of 0.05. Following the recommendations of Amrhein et al., we are reporting exact p-values and not classifying data into binary categories of ‘significant’ versus ‘insignificant’[43].

## Acknowledgements

Thank you to Nick Roma for his GC-MS work, and to Alexandra Domardsky and Sara Kass for their work on establishing FMPER protocols in the lab. Abigail Edwards, Katie Esbenshade, Vanessa Pham, Jessica Sommer, Baadal Vacchani, Katie Chen, Natalie David, Rachel Koerwer, Vijay Rao, and Morgan Tietz assisted in FMPER data collection. Many thanks to Allison Davidson for statistical consultation. Thank you to Muhlenberg College for research and travel funding.

## References

1. Joar Hegland S, Totland Ø. Is the magnitude of pollen limitation in a plant community affected by pollinator visitation and plant species specialisation levels? Oikos. Wiley Online Library; 2008;117: 883–891. doi:10.1111/j.2008.0030-1299.16561.x

2. Klein AM, Vaissiere BE, Cane JH, Steffan-Dewenter I, Cunningham SA, Kremen C, et al. Importance of pollinators in changing landscapes for world crops. Proc R Soc Biol. 2008;274: 303–313. doi:10.1126/science.265.5176.1170

3. Goulson D, Lye GC, Darvill B. Decline and conservation of bumble bees. Annu Rev Entomol. 2008;53: 191–208. doi:10.1146/annurev.ento.53.103106.093454

4. Whitehorn PR, O’Connor S, Wackers FL, Goulson D. Neonicotinoid pesticide reduces bumble bee colony growth and queen production. Science. 2012;336: 351–352. doi:10.1126/science.1215025

5. Rodríguez I, Gumbert A, Hempel de Ibarra N, Kunze J, Giurfa M. Symmetry is in the eye of the beeholder: innate preference for bilateral symmetry in flower-naïve bumblebees. Naturwissenschaften. Springer-Verlag; 2004;91: 374–377. doi:10.1007/s00114-004-0537-5

6. Odell E, Raguso RA, Jones KN. Bumblebee foraging responses to variation in floral scent and color in snapdragons (Antirrhinum: Scrophulariaceae). Am Midl Nat. 1999.

7. Leonard AS, Dornhaus A, Papaj DR. Flowers help bees cope with uncertainty: signal detection and the function of floral complexity. J Exp Biol. 2011;214: 113–121. doi:10.1242/jeb.047407

8. Sprayberry JDH. The prevalence of olfactory-versus visual-signal encounter by searching bumblebees. Sci Rep. Springer US; 2018;8: 1–10. doi:10.1038/s41598-018-32897-y

9. Spaethe J, Brockmann A, Halbig C, Tautz J. Size determines antennal sensitivity and behavioral threshold to odors in bumblebee workers. Naturwissenschaften. Springer; 2007;94: 733–739. doi:10.1007/s00114-007-0251-1

10. Sprayberry JDH, Ritter KA, Riffell JA. The effect of olfactory exposure to non-insecticidal agrochemicals on bumblebee foraging behavior. Frye MA, editor. PLoS ONE. 2013;8: e76273. doi:10.1371/journal.pone.0076273.s001

11. McFrederick QS, Kathilankal JC, Fuentes JD. Air pollution modifies floral scent trails. Atmos Environ. Elsevier; 2008;42: 2336–2348. doi:10.1016/j.atmosenv.2007.12.033

12. Fuentes JD, Chamecki M, Roulston T, Chen B, Pratt KR. Air pollutants degrade floral scents and increase insect foraging times. Atmos Environ. Elsevier Ltd; 9999;141: 361–374. doi:10.1016/j.atmosenv.2016.07.002

13. Girling RD, Lusebrink I, Farthing E, Newman TA, Poppy GM. Diesel exhaust rapidly degrades floral odours used by honeybees. Sci Rep. 2013;3. doi:10.1038/srep02779

14. Lusebrink I, Girling RD, Farthing E, Newman TA, Jackson CW, Poppy GM. The Effects of Diesel Exhaust Pollution on Floral Volatiles and the Consequences for Honey Bee Olfaction. Journal Of Chemical Ecology. 2nd ed. 2015;41: 904–912. doi:10.1007/s10886-015-0624-4

15. Leonard RJ, Vergoz V, Proschogo N, McArthur C, Hochuli DF. Petrol exhaust pollution impairs honey bee learning and memory. 2018;128: 264–273. doi:10.1111/oik.05405

16. Riffell JA, Lei H, Abrell L, Hildebrand JG. Neural Basis of a Pollinator’s Buffet: Olfactory Specialization and Learning in Manduca sexta. Science. 2013;339: 200–204. doi:10.1126/science.1225483

17. Locatelli FF, Fernandez PC, Smith BH. Learning about natural variation of odor mixtures enhances categorization in early olfactory processing. Journal Of Experimental Biology. 2016;219: 2752–2762. doi:10.1242/jeb.141465

18. Guerrieri F, Schubert M, Sandoz J-C, Giurfa M. Perceptual and Neural Olfactory Similarity in Honeybees. Plos Biol. 2005;3: e60. doi:10.1371/journal.pbio.0030060

19. Rering CC, Beck JJ, Hall GW, McCartney MM, Vannette RL. Nectar-inhabiting microorganisms influence nectar volatile composition and attractiveness to a generalist pollinator. New Phytol. John Wiley & Sons, Ltd (10.1111); 2017;220: 750–759. doi:10.1111/nph.14809

20. Hallem EA, Carlson JR. Coding of Odors by a Receptor Repertoire. 2006;125: 143–160. doi:10.1016/j.cell.2006.01.050

21. Clifford MR, Riffell JA. Mixture and odorant processing in the olfactory systems of insects: a comparative perspective. J Comp Physiol A. 2013;199: 911–928. doi:10.1007/s00359-013-0818-6

22. Sachse S, Rappert A, Galizia CG. The spatial representation of chemical structures in the antennal lobe of honeybees: steps towards the olfactory code. Eur J Neurosci. 1999;11: 3970–3982.

23. Galizia CG, Szyszka P. Olfactory coding in the insect brain: molecular receptive ranges, spatial and temporal coding. Entomol Exper Applic. 2008;128: 81–92. doi:10.1111/j.1570-7458.2007.00661.x

24. Shields VDC, Hildebrand JG. Responses of a population of antennal olfactory receptor cells in the female moth *Manduca sext*a to plant-associated volatile organic compounds. J Comp Physiol A. Springer-Verlag; 2001;186: 1135–1151. doi:10.1007/s003590000165

25. Carcaud J, Giurfa M, Sandoz J-C. Differential Processing by Two Olfactory Subsystems in the Honeybee Brain. Neuroscience. IBRO; 2018;374: 33–48. doi:10.1016/j.neuroscience.2018.01.029

26. Okada K, Kanzaki R. Localization of odor-induced oscillations in the bumblebee antennal lobe. Neurosci Lett. 2001;: 1–4.

27. Martin JP, Beyerlein A, Dacks AM, Reisenman CE, Riffell JA, Lei H, et al. The neurobiology of insect olfaction: Sensory processing in a comparative context. Progress in Neurobiology. 2011;95: 427–447. doi:10.1016/j.pneurobio.2011.09.007

28. Muth F, Cooper TR, Bonilla RF, Leonard AS. A novel protocol for studying bee cognition in the wild. Carvalheiro L, editor. Methods Ecol Evol. 2nd ed. 2017;9: 78–87. doi:10.1111/2041-210X.12852

29. Perez M, Giurfa M, d’Ettorre P. The scent of mixtures: rules of odour processing in ants. Sci Rep. 2015;5: 628–9. doi:10.1038/srep08659

30. Molet M, Chittka L, Raine NE. How floral odours are learned inside the bumblebee (Bombus terrestris) nest. Naturwissenschaften. 2008;96: 213–219. doi:10.1007/s00114-008-0465-x

31. Kulahci IG, Dornhaus A, Papaj DR. Multimodal signals enhance decision making in foraging bumble-bees. Proc R Soc Biol. The Royal Society; 2008;275: 797–802. doi:10.1098/rspb.2007.1176

32. Knauer AC, Schiestl FP. Bees use honest floral signals as indicators of reward when visiting flowers. Irwin R, editor. Ecol Letters. Wiley/Blackwell (10.1111); 2014;18: 135–143. doi:10.1111/ele.12386

33. Rusch C, Broadhead GT, Raguso RA, Riffell JA. Olfaction in context—sources of nuance in plant–pollinator communication. Current Opinion in Insect Science. Elsevier Inc; 2016;15: 53–60. doi:10.1016/j.cois.2016.03.007

34. Bos N, Dreier S, Jørgensen CG, Nielsen J, Guerrieri FJ, d’Ettorre P. Learning and perceptual similarity among cuticular hydrocarbons in ants. J Insect Physiol. Elsevier Ltd; 2012;58: 138–146. doi:10.1016/j.jinsphys.2011.10.010

35. Perez M, Nowotny T, d’Ettorre P, Giurfa M. Olfactory experience shapes the evaluation of odour similarity in ants: a behavioural and computational analysis. Proc Biol Sci. 2016;283: 20160551–9. doi:10.1098/rspb.2016.0551

36. Dornhaus A, Chittka L. Information flow and regulation of foraging activity in bumble bees (Bombusspp.). Apidologie. 2004;35: 183–192. doi:10.1051/apido:2004002

37. Granero AM, Sanz JMG, Gonzalez FJE, Vidal JLM, Dornhaus A, Ghani J, et al. Chemical compounds of the foraging recruitment pheromone in bumblebees. Naturwissenschaften. 2005;92: 371–374. doi:10.1007/s00114-005-0002-0

38. Hallem EA, Carlson JR. Coding of odors by a receptor repertoire. Cell. 2006;125: 143–160. doi:10.1016/j.cell.2006.01.050

39. Carcaud J, Hill T, Giurfa M, Sandoz JC. Differential coding by two olfactory subsystems in the honeybee brain. J Neurophysiol. 2012;108: 1106–1121. doi:10.1152/jn.01034.2011

40. Riffell JA, Shlizerman E, Sanders E, Abrell L, Medina B, Hinterwirth AJ, et al. Flower discrimination by pollinators in a dynamic chemical environment. Science. American Association for the Advancement of Science; 2014;344: 1515–1518. doi:10.1126/science.1251041

41. Riveros AJ, Gronenberg W. Olfactory learning and memory in the bumblebee Bombus occidentalis. Naturwissenschaften. Springer; 2009;96: 851–856. doi:10.1007/s00114-009-0532-y

42. McDonald JH. Handbook of Biological Statistics. 2009.

43. Amrhein V, Greenland S, McShane B. Scientists rise up against statistical significance. Nature. 2019.: 305–307. doi:10.1038/d41586-019-00857-9

