## Supplemental Methods and Tables for "Compounds without borders: a novel paradigm for quantifying complex odors and responses to scent-pollution in bumblebees"

Jordanna D. H. Sprayberry

**This PDF file includes:**

Supplementary Methods
Tables S1 to S3
Appendices S1 to S3

### Supplementary Methods

#### *Odor stimuli*

We selected three essential oils (*New Directions Aromatics)* to serve as odor-blends for this study: lily of the valley, honeysuckle and juniper berry. Lily of the valley (LoV) has been successfully used in odor learning experiments utilizing the proboscis extension reflex (PER) (41). The two additional odors were selected to provide a range of structural overlap with LoV: the floral-scent honeysuckle served as an ecologically similar odor; while the vegetative odor of juniper berry was selected as the 'dissimilar' odor. Coniferous vegetation is a pungent background in some bumblebee habitats; thus juniper berry represents an ecologically relevant odor pollutant. Blending these three in varying ratios allowed construction of odor stimuli with varying ranges of odorant composition and distance (Table 1). While these essential oils were selected for ecological relevance, commercial essential oils are not likely perfectly-accurate reproductions of their natural odor counterparts (Edwards et al., *in prep*). However, they are multi-component blends that will generate complex stimulation of the bumblebee olfactory system – likely pushing the system towards configural coding (42). These selected essential oils (EOs), as well as those from the predictive FMPER experiments, were sampled using Solid-Phase Microextraction (SPME) fibers and their composition analyzed with GCMS. For full details please see supplementary methods.

#### *Odor sampling and characterization of odor-blends*

Selected essential oils (EOs) were sampled using Solid-Phase Microextraction (SPME) fibers (Supelco 57359-U with PDMS/DVB coating) analyzed with a GC-MS. Thirty microliters of 1:1000 EO dilutions (in odorless mineral oil) were pipetted into glass sampling-vials fitted with a septum cap. The head-space within a vial was allowed to equilibrate at room temperature for 30 minutes, then the SPME fiber was injected and allowed to adsorb odorants for an additional 60 minutes. Fibers were then analyzed using a Shimadzu GC-QP5050A. Samples were injected into a splitless programmed-temperature injector to a Zebron ZB-5MS (5% phenyl, 95% diethylpolysiloxane) of 30 m x .2 mm i.d. x .25 µm film thickness analytical column from phenonenox. The GC temperature program was: an initial temp of 50 C, hold for 5 min, ramp at 10 c/min, finish at 320, hold for 15 min. Peaks on the GC spectra were identified through analysis of MS-spectra peaks with the connected chemical library; the compound with the highest similarity was chosen and all identified peaks had a similarity rating of 80% or above, with the majority of peaks (79%) having a similarity rating of 90% or higher. Once a putative molecular identity was assigned, peaks could be categorized in terms of functional group and carbon chain length. The normalized % area under identified peaks was calculated in LabSolutions.

#### *Statistical model of expected distribution for odor discrimination with FMPER*

Given that FMPER did indeed provide a reasonable measure of odor-learning (Fig. 3), the response-distribution of bumblebees tested with an AO vs MO should represent a simple odor discrimination task. We used data from the three AO vs MO experiments in this study, as well as four additional AO vs MO experiments from a separate methods study (Edwards et al. *in prep*) to establish the mean correct, incorrect and no-choice responses made by bumblebees in this simple task (Table S2). These data were used to statistically model the a theoretical response distribution of how bumblebees respond to an easy odor discrimination task in the FMPER assay (code available in Appendix S3). This model took a randomly generated number (from 0-1) and assigned it to one of five probability ranges derived from the overlap of the mean responses with their standard deviation (Fig. 2B): range 1 (0-0.56) had a 100% probability of being correct; range 2 (0.57-0.58) had an equal probability of being correct or no-choice; range 3 (0.59-0.72) had an equal probability of being correct, incorrect, or no-choice; range 4 (0.73-0.75) had an equal probability of being incorrect or no-choice; and range 5 (0.76-1) had a 100% probability of being no-choice. Once a randomly generated number, representing a silica-bee, was shunted into a range, its behavior was randomly selected from the possible outcomes. For example, bees assigned to range 1 (because the randomly-generated number was 0.23) would be categorized as correct; while a bee assigned to range 3 (because the randomly generated number was 0.65) would randomly be categorized as correct, incorrect, or no-choice. The model ran 100,000 silica-bees and returned a response distribution of 63% correct, 6% incorrect, and 31% no choice.

**Table S1. CWB dimensions are listed in the first column. The designation cyclicAlkene* is given to any molecules with carbon-carbon double bonds in a ring that are not aromatics. The designation monoterpene** is given to any monoterpenes or monoterpene derivatives. The light grey columns depict the CWB vectors for odorant-constituents of honeysuckle. These are then summed to create honeysuckle's CWB-vector. The juniper berry and lily of the valley vectors are also provided for comparative purposes.**

| | Honeysuckle components | | | | | | | | | Honey-suckle | Juniper Berry | Lily of the Valley |
|---|---|---|---|---|---|---|---|---|---|---|---|---|
| **CWB Dimensions** | octane | phenylethyl alcohol | Acetic acid, phenylmethyl ester | alpha-terpineol | butanoic acid | benzenepropanol | 2-propen-1-ol | benzoic acid | naphthalene | | | |
| acid | | | | | | | | | | 0 | 0.75 | 0.17 |
| acidChloride | | | | | | | | | | 0 | 0 | 0 |
| alcohol | | 26.9 | | 3.6 | | 4.7 | 4.8 | | | 40 | 23.53 | 82.61 |
| aldehyde | | | | | | | | | | 0 | 0 | 52.75 |
| alkane | 4 | | | | 4.1 | | | | | 8.1 | 8.2 | 2.95 |
| alkene | | | | | | | 4.8 | | | 4.8 | 46.46 | 15.81 |
| alkylHalide | | | | | | | | | | 0 | 0 | 0 |
| alkyne | | | | | | | | | | 0 | 0 | 0 |
| allylicMethyl | | | | | | | | | | 0 | 87.87 | 9.71 |
| amide | | | | | | | | | | 0 | 0 | 0 |
| amine | | | | | | | | | | 0 | 0 | 0 |
| aromatic | | 26.9 | 33.9 | | | 4.7 | | 7.8 | 10.2 | 83.5 | 0 | 29.29 |
| bicyclic | | | | | | | | | 10.2 | 10.2 | 46.33 | 0.6 |
| carboxylicAcid | | | | | 4.1 | | | 7.8 | | 11.9 | 0 | 0 |
| cyclic | | | | 3.6 | | | | | | 3.6 | 23.61 | 0.56 |
| cyclicAlkene* | | | | 3.6 | | | | | | 3.6 | 45.34 | 2.1 |
| epoxide | | | | | | | | | | 0 | 0 | 0 |
| ester | | | 33.9 | | | | | | | 33.9 | 10.22 | 6.4 |
| ether | | | | | | | | | | 0 | 0 | 0 |
| imide | | | | | | | | | | 0 | 0 | 0 |
| ketone | | | | | | | | | | 0 | 4.26 | 0.56 |
| lactone | | | | | | | | | | 0 | 0 | 0 |
| methyl | | | 33.9 | 3.6 | | | | | | 37.5 | 28.44 | 62.61 |
| monoterpene** | | | | 3.6 | | | | | | 3.6 | 80.73 | 58.05 |
| nitrile | | | | | | | | | | 0 | 0 | 0 |
| oxacyclic | | | | | | | | | | 0 | 0 | 0 |

| | | | | | | | | | | | | |
|---|---|---|---|---|---|---|---|---|---|---|---|---|
| oxabicyclic | | | | | | | | | | 0 | 0 | 0 |
| oxime | | | | | | | | | | 0 | 0 | 0 |
| phosphate | | | | | | | | | | 0 | 0.75 | 0.17 |
| thiol | | | | | | | | | | 0 | 0 | 0 |
| tricyclic | | | | | | | | | | 0 | 0 | 0 |
| CCL1 | | | | 3.6 | | | | | 10.2 | 13.8 | 0 | 0 |
| CCL2 | | | 33.9 | | | | | 7.8 | | 41.7 | 43.87 | 3.38 |
| CCL3 | | 26.9 | | | | | 4.8 | | | 31.7 | 26.07 | 28.84 |
| CCL4 | | | | | 4.1 | 4.7 | | | | 8.8 | 0 | 0.16 |
| CCL5 | | | | | | | | | | 0 | 0 | 0 |
| CCL6 | | | | | | | | | | 0 | 10.22 | 0 |
| CCL7 | | | | | | | | | | 0 | 0.75 | 0.17 |
| CCL8 | 4 | | | | | | | | | 4 | 19.09 | 67.45 |
| CCL9 | | | | | | | | | | 0 | 0 | 0 |
| CCL10 | | | | | | | | | | 0 | 0 | 0 |
| CCL11 | | | | | | | | | | 0 | 0 | 0 |
| CCL12 | | | | | | | | | | 0 | 0 | 0 |
| CCL13 | | | | | | | | | | 0 | 0 | 0 |
| CCL14 | | | | | | | | | | 0 | 0 | 0 |
| CCL15 | | | | | | | | | | 0 | 0 | 0 |
| CCC3 | | | | | | | | | | 0 | 0 | 0 |
| CCC4 | | | | | | | | | | 0 | 0 | 0 |
| CCC5 | | | | | | | | | | 0 | 0 | 0 |
| CCC6 | | 26.9 | 33.9 | 3.6 | | 4.7 | | 7.8 | | 76.9 | 26.07 | 31.78 |
| CCC7 | | | | | | | | | | 0 | 43.87 | 0.6 |
| CCC8 | | | | | | | | | | 0 | 0 | 0 |
| CCC9 | | | | | | | | | | 0 | 0 | 0 |
| CCC10 | | | | | | | | | 10.2 | 10.2 | 0 | 0 |
| CCC11 | | | | | | | | | | 0 | 0 | 0 |
| CCC12 | | | | | | | | | | 0 | 0 | 0 |
| CCC13 | | | | | | | | | | 0 | 0 | 0 |
| CCC14 | | | | | | | | | | 0 | 0 | 0 |

**Table S2. Data and analysis used to construct the probability model to generate an expected response distribution.**

| RAW DATA | | | | |
|---|---|---|---|---|
| Dataset | correct | incorrect | no response | p value (LLR) (compared to null) |
| LoV vs MO | 22 | 5 | 6 | 0.0006 |
| JB vs MO | 15 | 3 | 3 | 0.002 |
| Crd vs MO | 15 | 4 | 11 | 0.033 |
| PM vs MO | 15 | 0 | 5 | 0.000044 |
| SB vs MO | 16 | 0 | 13 | 0.000000088 |
| LD vs MO | 15 | 0 | 12 | 0.000036 |
| ME vsMO | 11 | 3 | 3 | 0.046 |
| **total bees used across experiments** | | | 177 | |
| **Response Ratio Calculations** | | | | |
| Dataset | correct | incorrect | no response | |
| LoV vs MO | 0.666666667 | 0.151515152 | 0.181818182 | |
| JB vs MO | 0.714285714 | 0.142857143 | 0.142857143 | |
| Crd vs MO | 0.5 | 0.133333333 | 0.366666667 | |
| PM vs MO | 0.75 | 0 | 0.25 | |
| SB vs MO | 0.551724138 | 0 | 0.448275862 | |
| LD vs MO | 0.555555556 | 0 | 0.444444444 | |
| ME vsMO | 0.647058824 | 0.176470588 | 0.176470588 | |
| **Descriptive Statistics** | | | | |
| mean | 0.626 | 0.086 | 0.287 | 1.00 |
| SD | 0.09 | 0.08 | 0.13 | |
| lower bound | 0.00 | 0.586 | 0.582 | |
| upper bound | 0.719 | 0.754 | 1.00 | |
| **Range Properties for Probability Model** | | | | |
| response | probability | lower | upper | |
| correct | 1 | 0 | 0.581 | |
| correct | 0.5 | 0.582 | 0.585 | |
| no response | 0.5 | 0.582 | 0.585 | |
| correct | 0.33 | 0.586 | 0.719 | |
| incorrect | 0.33 | 0.586 | 0.719 | |
| no response | 0.33 | 0.586 | 0.719 | |
| incorrect | 0.5 | 0.72 | 0.754 | |
| no response | 0.5 | 0.72 | 0.754 | |
| no response | 1 | 0.755 | 1 | |

**Table S3. Dimensional attributes of synthetic foraging pheromone constituents.**

| Compound | Carbon Chain Length | Cyclic Carbon Count | Functional Groups |
|---|---|---|---|
| Eucalyptol | 6 | 6 | oxabicyclic, methyl, monoterpene |
| ocimene | 8 | 0 | alkene, allylicMethyl |
| farnesol | 12 | 0 | alkene, alcohol, allylicMethyl |

**Appendix S1: R Code for CWB analysis**

```
#copy to .R file/s and adapt as necessary


##### first section is code to calculate vectors #####
#load dimensions
setwd("")
dimensions<- read.csv('dimensionMasterList.csv', header=F,stringsAsFactors =
FALSE) # you can create this from Table S1
unclass(dimensions)
#this has 3 categories of dimensions: functiontional group,  carbon chain length,
and cyclic carbons

#  load a .csv file of choice
odor_DATA <- read.csv("odorName.csv", header = T) #header should have
same format as odor tabs in supplemental dataset S1
#enter odor name
odorName = 'bob'
#master variable for the calculated vector
vectorCalc = matrix(data=NA,nrow=length(dimensions$V1),ncol=1)
#calculate vector signature
for(i in 1:length(dimensions$V1)){
  CCL<-sum(odor_DATA$area[which(odor_DATA$CCL==dimensions$V1[i])])
  CCC<-sum(odor_DATA$area[which(odor_DATA$CCC==dimensions$V1[i])])
  FG1<-sum(odor_DATA$area[which(odor_DATA$FG1==dimensions$V1[i])])
  FG2<-sum(odor_DATA$area[which(odor_DATA$FG2==dimensions$V1[i])])
  FG3<-sum(odor_DATA$area[which(odor_DATA$FG3==dimensions$V1[i])])
  FG4<-sum(odor_DATA$area[which(odor_DATA$FG4==dimensions$V1[i])])
  FG5<-sum(odor_DATA$area[which(odor_DATA$FG5==dimensions$V1[i])])
  FG6<-sum(odor_DATA$area[which(odor_DATA$FG6==dimensions$V1[i])])
  vectorCalc[i]<-sum(CCL,CCC,FG1,FG2,FG3,FG4,FG5,FG6)
}

#build vectors into a .csv file

##### the second section is code to calculate angles #####

#load file
setwd("")
d<- read.csv('myVectors.csv', header=T,stringsAsFactors = FALSE)
d2<-d[,-1] #drop first column of d in d2
rownames(d2)<-d[,1] #use that dropped column as names
attach(d2)
```

```
#Which vectors are we calculating the angle between?
angleName = 'bobToBetty'
v1 = bob # name of an odor vector
v2= betty # name of an odor vector
#calculate vector length for all odors
a=v1*v1
vectorLength1 = sqrt(sum(a))
b=v2*v2
vectorLength2 = sqrt(sum(b))
#Dot Products
dotVectors = v1%*%v2
#angle calculation
Theta=acos(dotVectors/(vectorLength1*vectorLength2))
#assemble angle entry
angleData<-c(angleName,Theta)
```

### Appendix S2: FMPER Protocol

This protocol is adapted from:

F. Muth, T. R. Cooper, R. F. Bonilla, A. S. Leonard, A novel protocol for studying bee cognition in the wild. *Methods Ecol Evol*. **9**, 78–87 (2017).

#### Experimenter Preparation

- Be aware of potential scent contamination - these are odor learning experiments and we want control over what odors bees are exposed to
- glove early, glove often. You should ALWAYS wear gloves when handling bees or experimental materials to prevent your skin oils/ lotions/ etc from leaving a scent residue

#### Bee Preparation

- Collect candidate bees and place them in the modified tubes. Allow bees to acclimate to tubes for 2-3 hours - if you collect 9-12 bees you will likely be running active tests for approximately an hour. The FMPER ventilation array holds 6 bees at a time, so this is the recommended minimum number to test. If you do test less you will need to insert empty tubes into the array to assure that airflow is consistent across trials.

#### Stimulus strip preparation

- Prior to experiments set up your conditioning and testing strips (this typically takes 15-20 minutes prior to actual testing)
- Make plastic strips by cutting an 1.75"-2" wide rectangle from a yellow plastic folder. Then cut a piece of 0.5"-0.75" wide stretch adhesive bandage cover and affix it to the long end of your plastic rectangle. At this point you should be ready to cut thin plastic strips for use in PER experiments. The strips will have one side that is entirely plastic, and one side that has the lower half covered with absorbent bandage tape. The tape is difficult to affix to individual thin strips, which is why you make a large block prior to cutting strips. Store any extra strips you made in a clean vial for future experiments.
- Take out three small weigh boats: one for your AO+, one for your AO-, and one for your CO-. Add two strips to the AO+ and one to the AO- and the CO-. Fllip the strips so that the absorbent tape is facing up.
    - AO = your associative odor
    - CO = your contrasting odor
    - + indicates a reward will be present
    - - indicated no reward
- In the hood pipette 1 $\mu$L of the AO onto all three AO strips. Then pipette 1uL of the CO onto the CO strip. If the CO is a blend, you can serially apply the relevant volumes to the tape. Be sure to switch tips between odors to prevent contamination of the stocks. Once you are finished, change your gloves to reduce potential odor contamination

#### Associative Conditioning Phase

- Set out a small (10 ml) beaker of 50% sucrose solution with a toothpick next to the FMPER rig. Using chalk or a chalk marker, box out and label the locations for your AO+, AO-, and CO- strips. Make sure you have a timer, a data collection board, a die, and a whiteboard marker
- Put 6 of your acclimated bees into the FMPER rig. Using the toothpick place a drop of sucrose on an AO+ strip. Then introduce the strip into a vial to entice a bee for a snack. ONE TIME you can tap the sugar side of the strip on the floor of the vial and leave a drop of sugar behind. This can serve to prime bees and make them more willing to participate. The reasoning behind only doing this once is that you want to build an association between the scented strip and the reward, so

once conditioning has started they should only receive the reward from the strip.

- Once a bee has drunk from the strip, they have entered conditioning. Mark this as their first trial on the data collection board, being sure to note the side the strip was introduced on (from experimenter perspective, not the bee's). If this is the first bee to participate, start a five minute timer. If it is not, mark the time on the timer, you will continue to test at that point in the five minute cycle from here on out.
- For bees in the conditioning phase: every five minutes introduce the AO+ with a fresh drop of sucrose. Use the die to randomize which side the strip is introduced through (I use rolls ≤ 3 to indicate left, ≥ 4 to indicate right). If this is the fourth conditioning trial and the previous three have all been the same side, don't randomize - just introduce on the opposite side (i.e. if they have had R, R, R for the first three insert the strip on the L). If they do not drink immediately give them about 45 seconds before marking them no response. If they are no response, do not continue to condition them.
    - data entry in this phase: mark the side the strip was entered through ("L" or "R") and mark response as "Y" if they drink, "N" if they do not.
- Once bees have had 4 conditioning trials they move into the testing phase
- If you have tried to entice a bee for more than fifteen minutes with no success and you have additional acclimated bees waiting in the wings you can swap them out. DO NOT remove them if you don't have extra bees, PROPER AIR FLOW REQUIRES THERE TO BE 6 VIALS IN THE RIG FOR ALL EXPERIMENTS. Sorry about the all caps - if for some reason you do not have all six ports with a vial, make a note in the notes section of the data entry widget and we will discard the data during analysis. Remember, mistakes happen. What we don't want is for mistakes to contaminate data analysis, so be sure to note any issues in that notes section.
- Things to look out for:
    - don't let sucrose build up around holes in lid, use a kimwipe or q-tip to keep it clean
    - if sucrose gets smeared on strip, clean it
    - if odor tape on AO+ strip starts to pull off, switch to your extra (this is why I recommend prepping one)
    - if you are doing a large number of experiments, switch to your fresh strip after bee 9 or so

**Testing Phase**

- Use the die to determine which side the AO will be inserted on
- Get the AO- and CO- ready to insert on their proper sides
- Wait for the bee to be in the back half of the vial
- Insert strips simultaneously
- Watch the bee's behavior. If the bee extends it proboscis onto a strip, or while actively touching a strip with its antenna that counts as a choice.
    - if proboscis is extended on the AO- strip, mark as "C" for correct
    - if proboscis is extended on the CO- strip, mark as "I" for incorrect
- If bee does not choose immediately continue to watch behavior. Allow them to interact with the strips. Take note of when they disengage and walk away. Allow them to interact up to THREE times - after this you remove the strips and mark them "NR" for no response/ no choice.
    - To date we have not had a bee that didn't approach or interact with the strips during testing phase, if you encounter this you would leave the 'test response' section of the data widget on 'make choice' and write it up in the notes section. These data could then be discarded during the analysis phase.

**Post experiment breakdown**

- All bees that entered conditioning phase should get put on ice to immobilize/ slow them for marking. Bees that never participated at all can be put back in colony unmarked.
- To mark bees:
    - ice until calm
    - get out the super glue gel, forceps and glitter (I recommend making a little clay snake on the counter and using the forceps to stick glitter it, so that the glitter is sticking out and easy to pick up when you are trying to attach it to a bee)
    - once bees are chill (get it?), put them on the counter, put a dab of super glue on their thorax above the wing attachment, then place glitter on the glue blob. Tap the glitter down with the forceps to embed it in the glue. Put a screen-backed vial upside down over bee and let the glue dry/ bee wake up.
    - Once bees are moving and the glue is dry, reintroduce them into the colony. Put them back into the colony box proper, not the foraging chamber (this seems to reduce rejection of marked bees).
- Wash all vials in hot water and soap. Leave overnight for scent marks and odor residues that somehow made it through washing (they shouldn't) to dissipate
- discard strips and weigh boats
- wash sucrose beaker
- ENTER YOUR DATA IN THE WIDGET ON LAB ARCHIVES. Otherwise, what is the point in doing the experiments?

**Appendix S3: R Code for generating expected response distribution**

```
#copy to .R file, amend with relevant data/ range boundaries

# using experimental data to create a theoretical response distribution
# for an "easily discriminated" odor in FMPER
# See Table S2 for underlying data
# Three response possibilities: correct, incorrect, no choice
# Datasets for seven different odor stimuli tested against unscented mineral oil
# yielded the following averages/SDs for each response possibility
# (one data that differed significantly from a random null distribution were
included)
#          mean/SD
# Correct: 0.626/.09
# Incorrect: 0.086/0.08
# No choice: 0.287/0.13

# Approach: set up ranges from 0-1 with overlapping
# probability of an individual bee responding C, I, NC - these ranges
# have 1 SD range for each response, such that
# range 1: 0-0.581 has a 100% probability of yielding C
# range 2: 0.582-0.585 has an equal probability of yielding C or NC
# range 3: 0.586-0.719 has an equal probability of yielding C, I, or NC
# range 4: 0.72-0.754 has an equal probability of yielding I or NC
# range 5: 0.755-1 has a 100% probability of yielding NC

n = 100000 # number of simulated bees
Ns=runif(n)
results = matrix(data=NA,nrow=length(Ns),ncol=1)
range1Min = 0.00000
range1Max = 0.58199
r1=which(Ns>=range1Min & Ns<=range1Max)
results[r1]='C'

range2Min = 0.58200
range2Max = 0.58599
r2=which(Ns>=range2Min & Ns<=range2Max)
r2Responses=c('C','NC')
r2Results = sample(r2Responses,length(r2),replace=TRUE)
results[r2]=r2Results

range3Min = 0.58600
range3Max = 0.71999
```

```
r3=which(Ns>=range3Min & Ns<=range3Max)
r3Responses=c('C','I','NC')
r3Results = sample(r3Responses,length(r3),replace=TRUE)
results[r3]=r3Results

range4Min = 0.7200
range4Max = 0.75499
r4=which(Ns>=range4Min & Ns<=range4Max)
r4Responses=c('I','NC')
r4Results = sample(r4Responses,length(r4),replace=TRUE)
results[r4]=r4Results

range5Min = 0.75500
range5Max = 1
r5=which(Ns>=range5Min & Ns<=range5Max)
results[r5]='NC'

percentCorrect = length(which(results=='C'))/length(results)
percentIncorrect = length(which(results=='I'))/length(results)
percentNoChoice = length(which(results=='NC'))/length(results)

expectedEasyDiscrimination =
c(percentCorrect,percentIncorrect,percentNoChoice)

# after 100,000 iterations, program returned:
#   expectedEOC = c( 0.62750, 0.06101, 0.30767), round to 63%, 6%, 31%
```

**Additional data table S1 (separate file)**
All data for CWB analysis of test odors available for download as a .xls file.

**Additional data table S1 (separate file)**
All data from the presented FMPER experiments are available for download as a .xls file.
